# Predictive Representations in Hippocampal and Prefrontal Hierarchies

**DOI:** 10.1101/786434

**Authors:** Iva K. Brunec, Ida Momennejad

## Abstract

As we navigate the world, we use learned representations of relational structures to explore and to reach goals. Studies of how relational knowledge enables inference and planning are typically conducted in controlled small-scale settings. It remains unclear, however, how people use stored knowledge in continuously unfolding navigation, e.g., walking long distances in a city. We hypothesized that multiscale predictive representations guide naturalistic navigation, and these scales are organized along posterior-anterior prefrontal and hippocampal hierarchies. We conducted model-based representational similarity analyses of neuroimaging data measured during navigation of realistically long paths in virtual reality. We tested the pattern similarity of each point–along each path–to a weighted sum of its successor points within predictive horizons of different scales. We found that anterior PFC showed the largest predictive horizons, posterior hippocampus the smallest, with the anterior hippocampus and orbitofrontal regions in between. Our findings offer novel insights into how cognitive maps support hierarchical planning at multiple scales.

## Introduction

When we navigate a city we draw on our memory to get around. The structure of learned representations retrieved from memory reflects the multiscale structure of our environment. Relational structures (e.g., directions and distances between familiar locations) are learned, retrieved, and updated using such representations. This relational knowledge is later retrieved to make decisions, plan, and guide behavior (Behrens et al., 2018; Momennejad, 2020). This idea has been captured by computational models that account for planning in human behavior (Momennejad et al., 2017), relational knowledge in human functional magnetic resonance imaging (fMRI) (Garvert et al., 2017), and place and grid fields in rodent electrophysiology (Stachenfeld et al., 2017). This converging body of evidence suggests that the brain learns predictive representations of relational structures, which can be used for fast and flexible planning. It has been suggested that these predictive representations are organized in a multiscale fashion, each scale of representation corresponding to different gradients in the neural representational hierarchy, e.g., in the hippocampus (Momennejad & Howard, 2018; Stachenfeld et al., 2017) and the prefrontal cortex (Christoff & Gabrieli, 2000; Koechlin & Hyafil, 2007; Momennejad & Haynes, 2013).

Here, we tested the hypothesis that predictive representations are organized along prefrontal and hippocampal hierarchies at multiple scales, consistent with predictions made by our computational models. We predicted that representations at multiple scales would be active simultaneously, and supported by different brain regions. Such hierarchical structure in the representations of states and trajectories could also enable the extraction of generalized schemas, or structured relationships at higher levels of abstraction that can be unfolded at lower levels when necessary (Figure 1A).

**Figure 1.**
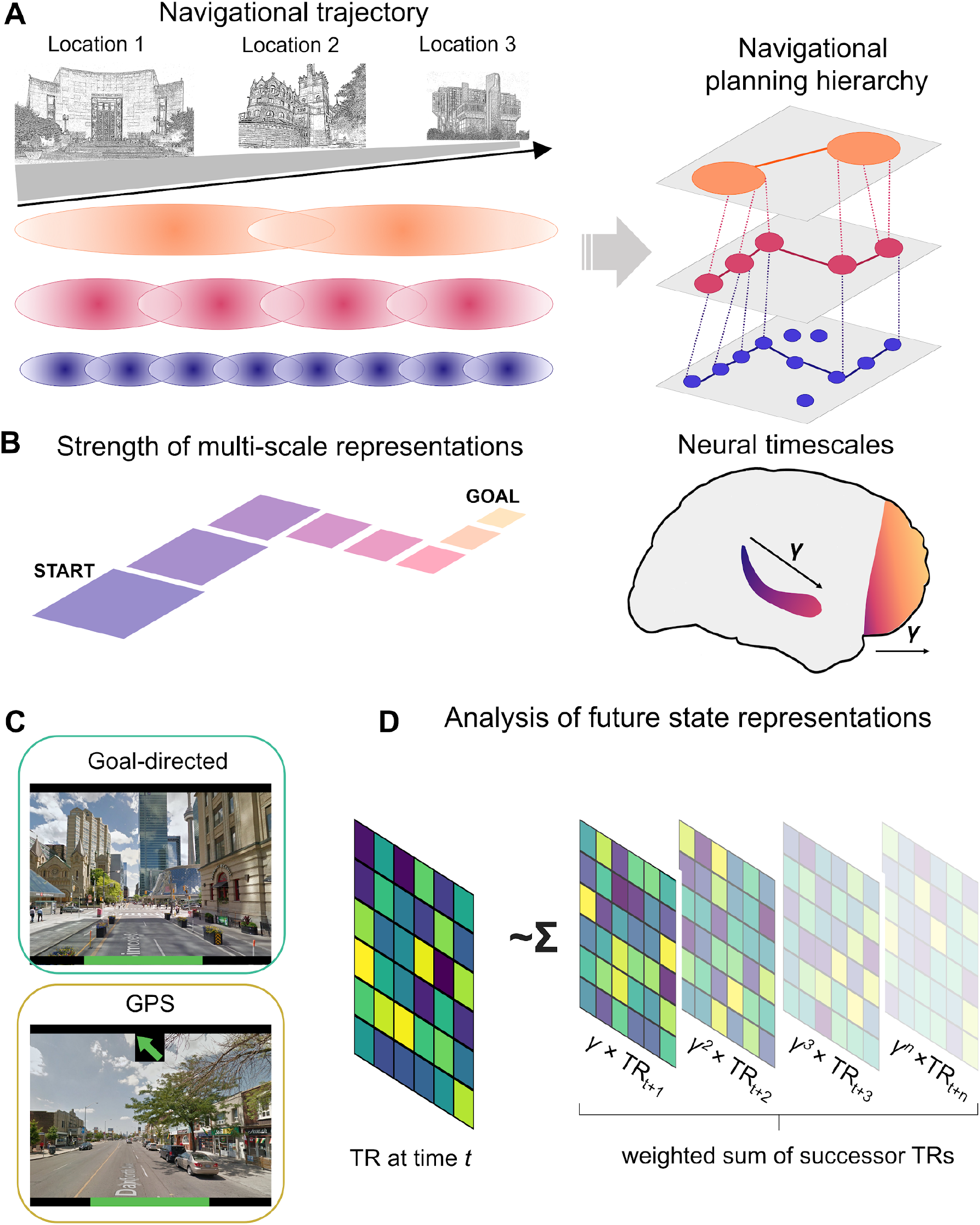
Schematic of the hypothesis, task conditions, and analytic methods. A) Multiple scales of representation along a navigated route are activated simultaneously. Longer predictive horizons correspond to longer-range planning and greater scales of navigational representations. B) Predictive representations in the hippocampus and prefrontal cortex should proceed along a posterior-anterior gradient within the hippocampus and prefrontal cortex. C) Participants used the same keys to navigate Goal-directed routes and to follow the GPS dynamic arrow, but only Goal-directed routes required goal-directed navigation. D) Analytic approach. The voxelwise pattern at each timepoint was correlated with the gamma-weighted sum of all future states (for gamma values of .1, .6, .8 and .9). With higher gamma values, the weighted future states remain above zero further into the future. Not displayed: We also computed similarity for each TR to goal, and similarity of each TR to mean of future TRs (equally weighted) within a given horizon (e.g. 10 TRs).

**Figure 2.**
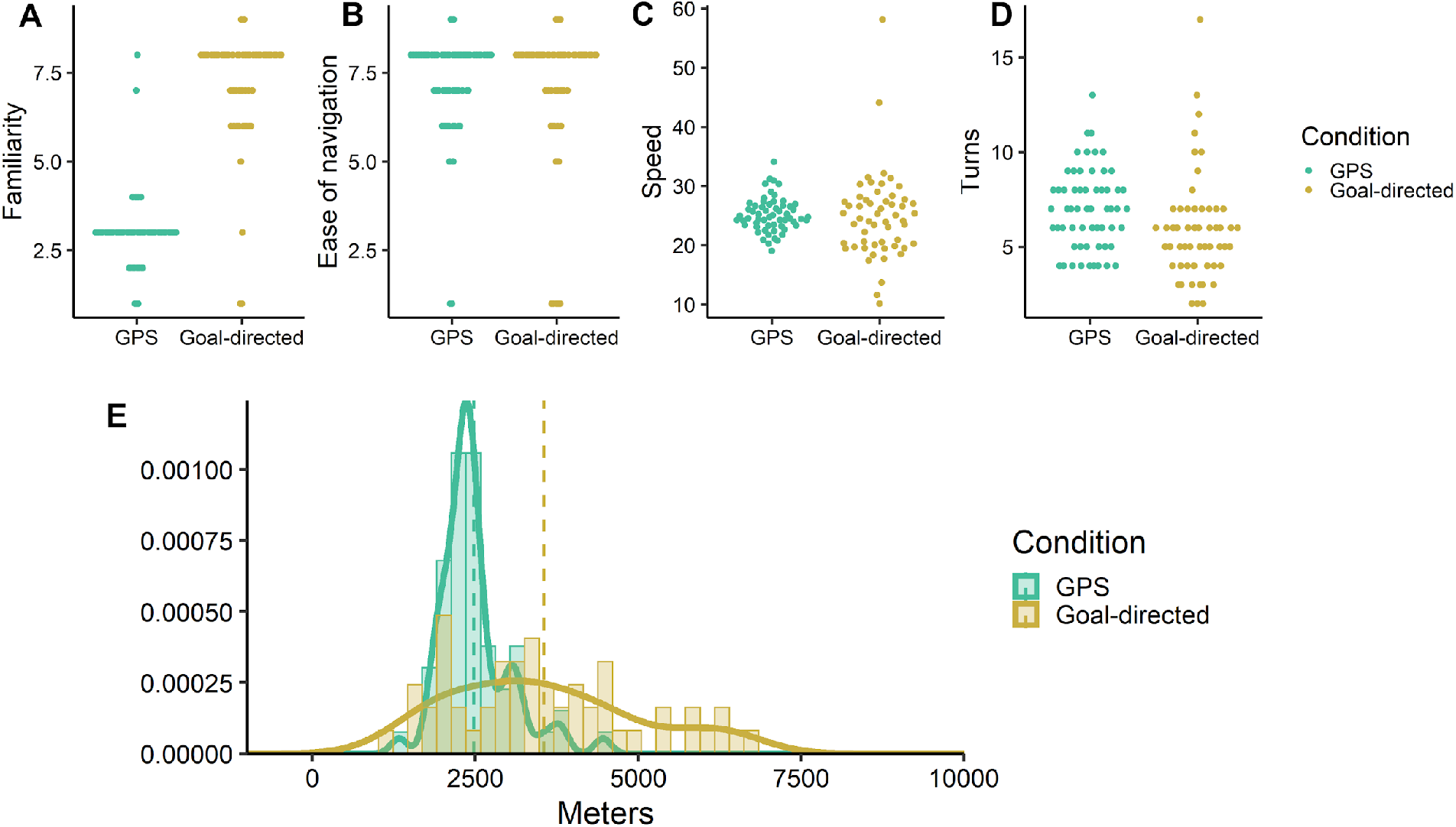
Descriptive statistics for navigated distances in Goal-directed and GPS conditions. A) The Goal-directed routes were rated as more familiar by participants than GPS routes. Goal-directed and GPS routes were matched in B) ease of navigation and C) speed of travel. D) GPS routes included more turns, on average, than Goal-directed routes, but E) the Goal-directed routes tended to be longer than GPS routes.

**Figure 3.**
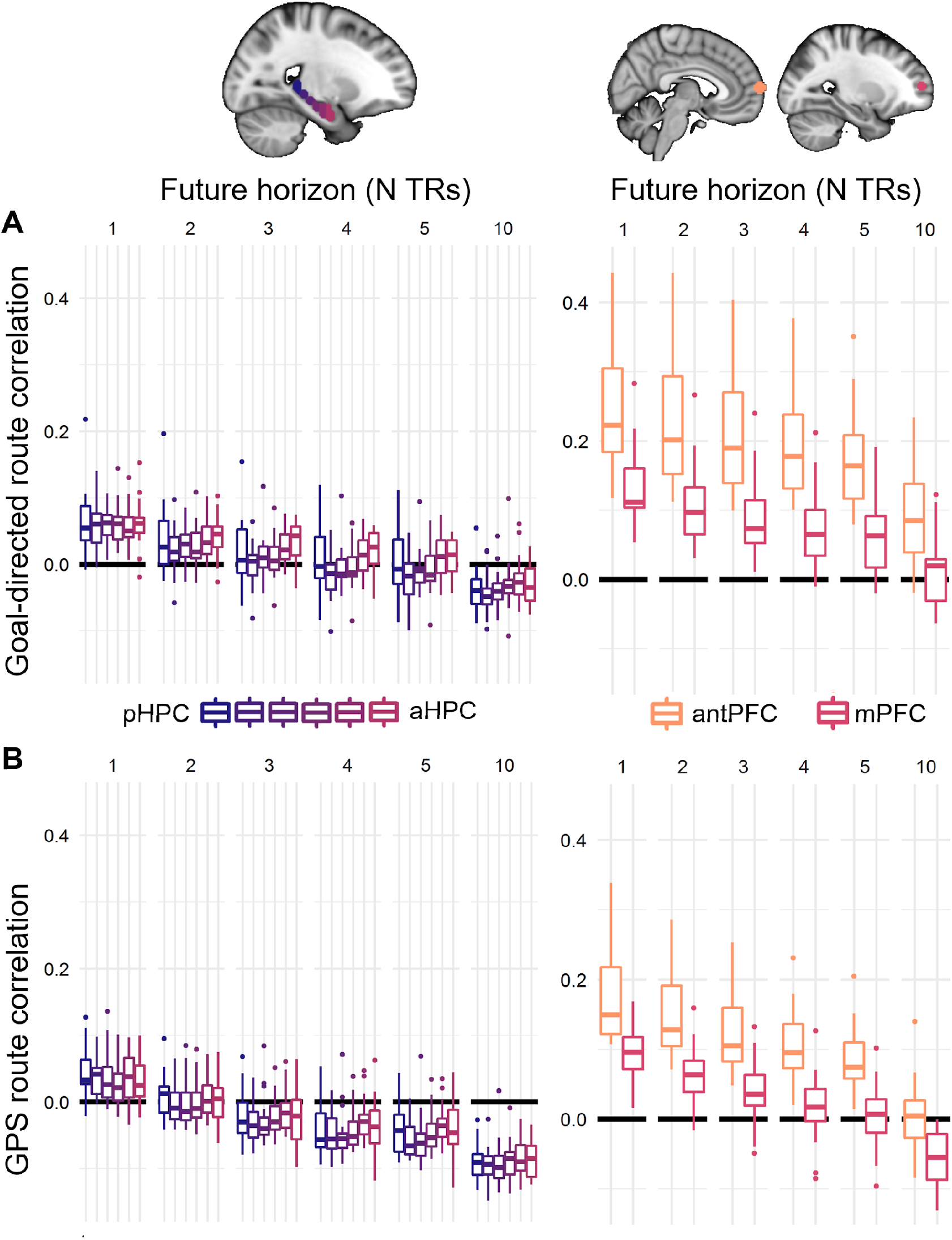
Similarity of each TR to mean of future TRs (equally weighted). Average correlation between each timepoint and the average of future 1/2/3/4/5 or 10 timepoints in A) the Goal-directed condition and B) the GPS condition. As the average distance traversed within each TR was 25 m (meters).

We hypothesized that simple principles of representation learning from our computational models may predict representations in human brains during navigation in everyday settings. Typically computational models are only tested in small-scale and highly controlled experiments. We used model-based analysis of fMRI data to test predictions of a reinforcement learning (RL) model (Momennejad & Howard, 2018) on brain signals collected during virtual navigation of a familiar real-world city, i.e. Toronto. We show that even though participants had learned these paths in their real lives, representations in their prefrontal and hippocampal hierarchies followed representations predicted by our hypothesized RL model. Previously we had shown that the representations learned by these models capture human behavior in planning tasks (Momennejad et al. 2017). We think this contribution opens up a new window into building and testing neurally plausible models of everyday cognition. Such models need to capture both behavioral and neural responses of humans performing the given task. Our approach offers a theory-rich perspective on testing models of multiscale predictive representations in the hippocampus and the PFC using fMRI representational similarity analysis.

Our framework of multiscale representations is grounded in decades of empirical findings from electrophysiology and neuroimaging of rodent and human brains. Hippocampal place cells fire within spatial fields of different sizes, and entorhinal grid fields tile the space at various levels of granularity (Brun et al., 2008; Kjelstrup et al., 2008; Poppenk et al., 2013; Strange et al., 2014). Evidence from rodent electrophysiology suggests that the average place field size increases along the dorsoventral axis of the rodent hippocampus, with more ventral regions encoding space at a larger spatial scale and in a more overlapping manner (Contreras et al., 2018; Jung et al., 1994; Strange et al., 2014). Furthermore, human fMRI evidence suggests that the hippocampal posterior-anterior axis (homologous to the rodent dorsal-ventral axis) is also involved in finer- to coarser-grained spatial representations (Evensmoen et al., 2013), and in inference from lower to higher levels of abstraction (Collin et al., 2015). Furthermore, the anterior hippocampus has been found to represent realistically long temporal distances between everyday events in fMRI (Nielson, Smith, Sreekumar, Dennis, & Sederberg, 2015).

Our first hypothesis concerned the role of the anterior-posterior hippocampus in multiscale predictive representation. The larger scale representations in the anterior hippocampus are proposed to support goal-directed search (Ruediger et al., 2012), the integration of spatial and non-spatial states that are further apart (Collin et al., 2015), and longer time horizons (Brunec, Bellana et al., 2018; Nielson, Smith, Sreekumar, Dennis, & Sederberg, 2015). The representations in the posterior hippocampus are more myopic and might support smaller predictive scales, such as smaller place fields in spatial navigation (Strange et al., 2014) and more pattern separation in memory studies (Duncan & Schlichting, 2018; Leutgeb et al., 2007; Lohnas et al., 2018; Schlichting et al., 2015). Recent computational models provide further support for multiscale predictive cognitive maps. These models account for why place fields are skewed toward goal locations (Stachenfeld et al., 2017) and show that multiscale predictive maps can capture distance to goal and reconstruct predicted sequences (Momennejad & Howard, 2018). Grounded in these findings, we hypothesized that in virtual navigation anterior hippocampus would display representational similarity at longer predictive scales compared to posterior hippocampus.

Our second hypothesis concerned the role of the anterior-posterior prefrontal cortex (PFC) in multiscale predictive representation. Another candidate region for processing hierarchical representations during planning is the prefrontal cortex (Badre & D’Esposito, 2007). Broadly, it has been proposed that the PFC is involved in navigation when it is *active* and requires planning (Behrens et al., 2018; Epstein et al., 2017; Spiers & Gilbert, 2015), in consideration of the number of paths to goal and alternative paths (Javadi et al., 2017), in reversal and detours (Spiers & Gilbert, 2015), and in retrospective revaluation mediated by offline replay (Momennejad, Otto, Daw, & Norman, 2018). Neuroimaging evidence suggests a prefrontal hierarchy in which more anterior PFC regions support relational reasoning (Christoff & Gabrieli, 2000; Christoff, Keramatian, Gordon, Smith, & Mädler, 2009), abstraction (Bunge, Kahn, Wallis, Miller, & Wagner, 2003; Christoff et al., 2001), and prospective memory (Gilbert, 2011; Haynes & Rees, 2006; Momennejad & Haynes, 2012, 2013). Bringing these findings together, we hypothesized that anterior PFC would display representational similarity to more distant states (representing longer predictive horizons, or larger scales), while more posterior PFC regions would display representational similarity within more myopic scales (representing shorter predictive horizons, or smaller scales).

Comparing the PFC and the hippocampus, we hypothesized that the scales of anterior PFC representations would exceed the longest predictive horizons of hippocampal representations (Figure 1B). PFC cells broadly have longer delays and more recurrent interconnectivity, enabling information to linger across longer scales and learning to occur at slower rates. The anterior PFC is the largest cytoarchitectonic region of the human prefrontal cortex, and a region in which we differ the most with evolutionary ancestors (Ramnani & Owen, 2004). In contrast, the hippocampus has been suggested to be involved in rapid statistical learning at a faster rate (Schapiro et al., 2017) and is less heterogeneous across mammal species (Strange et al., 2014).

To test the hypotheses of multiscale hippocampal-prefrontal hierarchies, we conducted model-based representational similarity analyses (Figure 1D) on an existing fMRI dataset of realistic virtual navigation (Brunec, Bellana et al., 2018). In this paradigm, participants navigated Goal-directed and GPS-guided routes in a virtual version of the city they lived in (Toronto). Virtual Toronto was built using images from Google Street View. In the Goal-directed condition, participants navigated routes that they regularly traversed in their everyday lives. In the GPS-guided condition, they were guided along novel routes by following a dynamic arrow. This virtual navigation setup had important advantages for our purposes. The participants’ experience was as realistic as possible within the constraints of fMRI, with everyday navigation at the scale of kilometers. Furthermore, the experimental design benefited from participants’ real world familiarity. This allowed us to compare the scales or predictive horizons of well-learnt long routes vs. novel routes. To navigate GPS-guided routes, participants used the same control buttons as they did along Goal-directed routes. However, in the GPS condition they did not know the goal or the distance (Figure 1C).

Consistent with our prediction, we found that on Goal-directed, compared to GPS-directed paths, anterior hippocampal and anterior PFC regions displayed representational similarity at longer horizons (i.e. displayed similarity to more distal states), while the posterior hippocampus supported predictive representations at smaller scales (i.e. displayed similarity to more proximal states). Notably, anterior PFC showed the longest predictive horizons (Figure 4).

**Figure 4.**
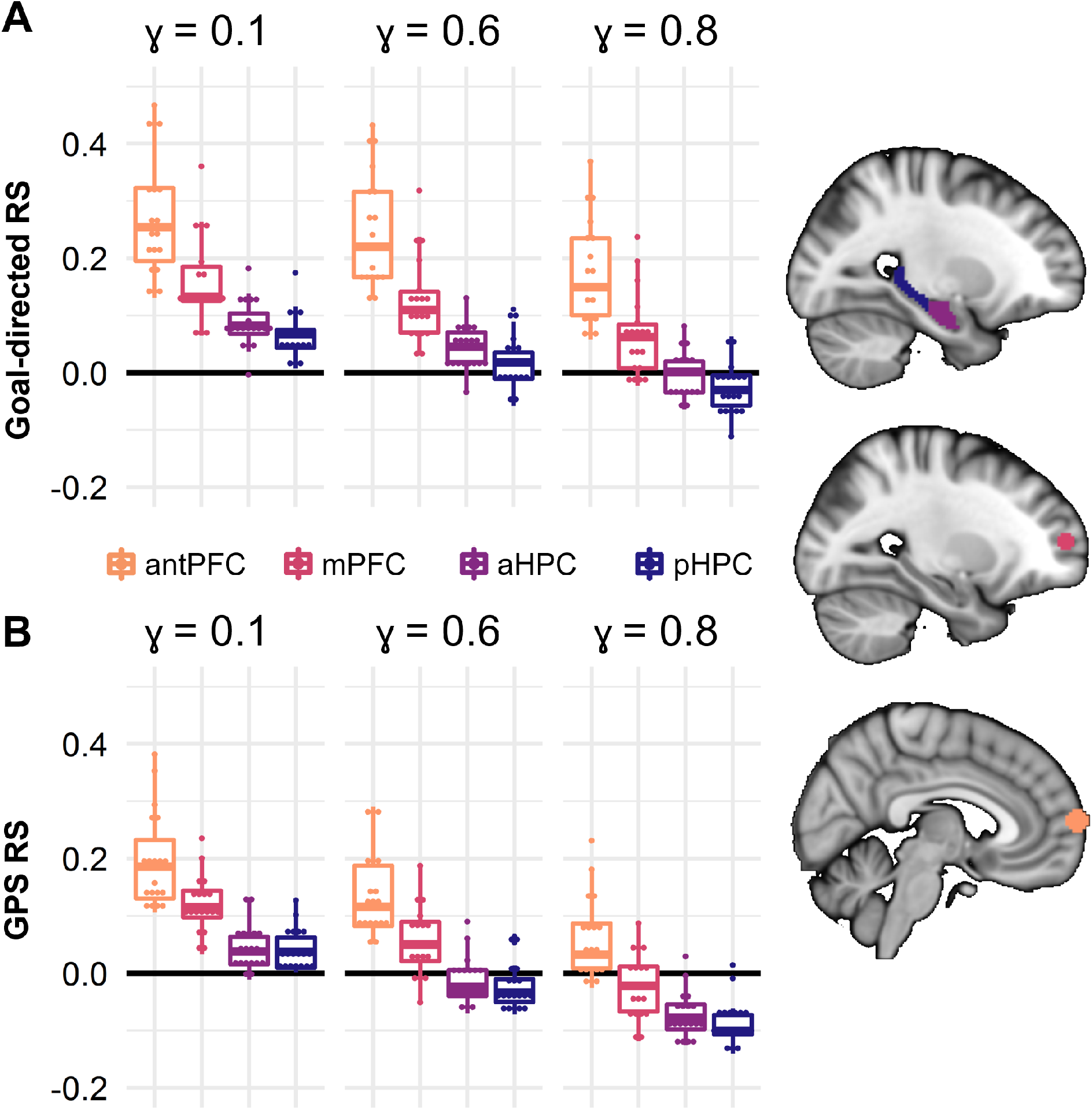
Predictive similarity across predictive scales. Correlations between current timepoints and the ɣ-weighted sum of future states for different values of gamma, in the four specified ROIs in the A) Goal-directed and B) GPS conditions. ɣ=.1 only included 1-step (1 TR) away, ɣ=.6 reached approximately 6-7 steps in the future, corresponding to roughly 175 m, ɣ=.8, approximately 14 steps or 350 m ahead.

**Figure 5.**
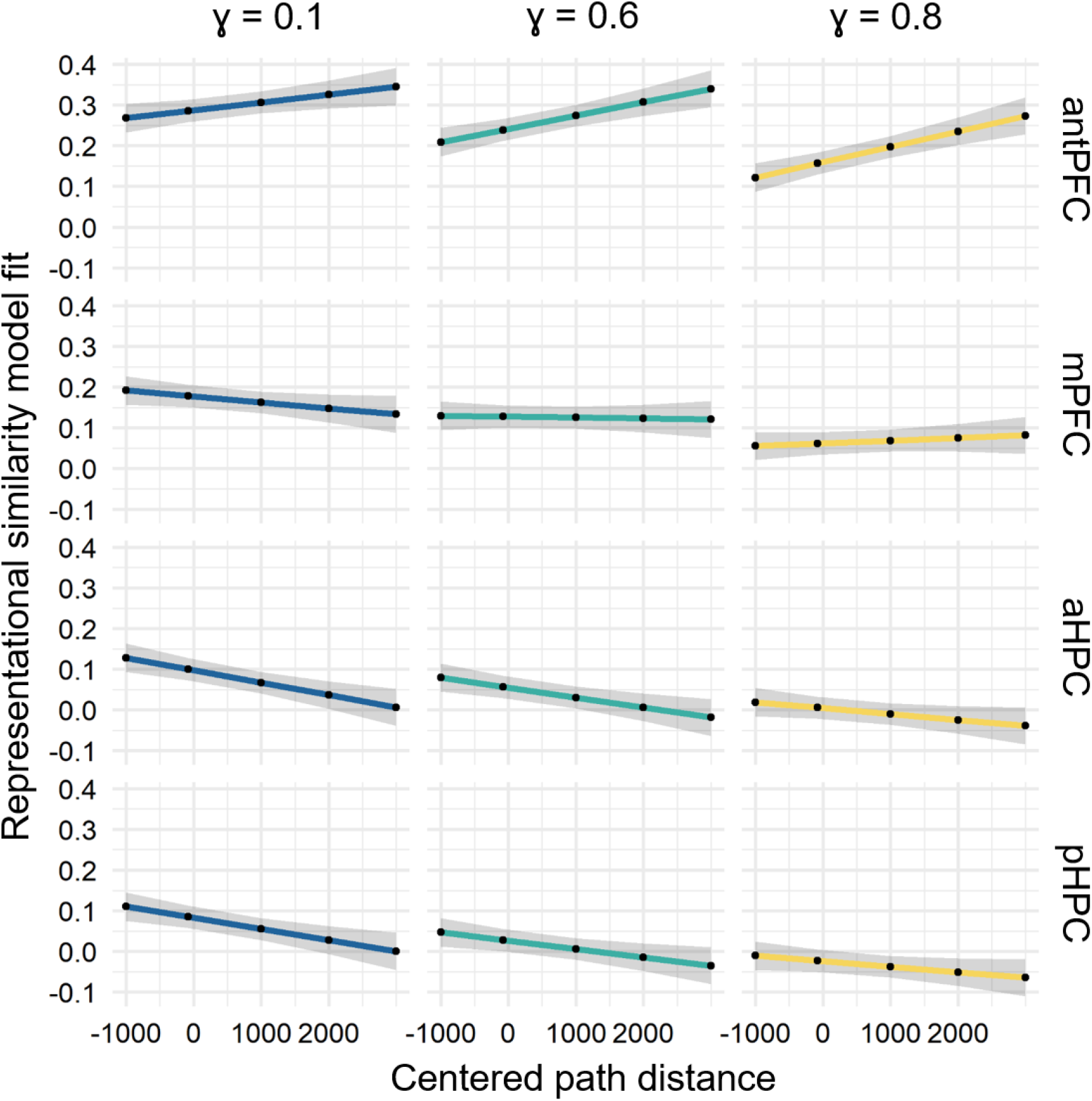
Linear mixed effects model predicting representational similarity (y-axis) from path distance (x-axis), ɣ, and ROI. Voxelwise patterns in different ROIs interacted differently with path distance: in the antPFC, routes with longer path distances were associated with greater representational similarity, while the opposite trend was present in the hippocampus (both aHPC and pHPC). The plot depicts the model fit values and confidence intervals. These reflect the relationships between the variables of interest after regressing out the effect of the number of TRs on each route and accounting for all other main effects and interactions.

## Methods

### Subjects

Twenty-two healthy right-handed volunteers were recruited. One participant was excluded because of excessive difficulty with the task (i.e., repeatedly getting lost). Two additional participants were excluded due to incomplete data or technical issues. Exclusions resulted in 19 participants who completed the study (9 males; mean age 22.58 years, range 19-30 years). All participants had lived in Toronto for at least 2 years (M = 10.45, SE = 1.81). All participants were free of psychiatric and neurological conditions. All participants had normal or corrected-to-normal vision and otherwise met the criteria for participation in fMRI studies. Informed consent was obtained from all participants in accordance with Rotman Research Institute at Baycrest’s ethical guidelines. Participants received monetary compensation upon completion of the study.

### Experimental design and paradigm

We used a realistic navigation software drawing on 360° panoramic images from Google Street View. This allowed participants to walk through a virtual Toronto from a first-person, street-level perspective. The navigation software was written in MATLAB v7.5.0.342. Navigation was controlled using three buttons: left, right, and forward. A “done” button allowed participants to indicate that they had completed a route. The task was projected on a screen in the bore of the scanner viewed by the participants through a mirror mounted inside of the head coil. Participants navigated in 4 conditions, and navigated 16 routes in total (4 in each condition, in a randomized order). The details of the experimental design have been reported in a previously published study (Brunec, Bellana, et al., 2018).

Data from two conditions of interest were analyzed in the present manuscript: Goal-directed and GPS/arrow-following routes. The routes were constructed prior to the day of scanning: participants built routes with researcher assistance, using a computer program which showed overhead maps of Toronto. Additionally, sets of routes in areas of Toronto with which participants were generally unfamiliar were created. Four of these routes were randomly assigned to each participant to be used in the baseline (GPS) condition. In the scanner, participants were provided with Goal-directed route destinations and asked to navigate towards the goal along the most Goal-directed/comfortable route. GPS trials involved no goal-directed navigation; instead, participants followed a dynamic arrow (Figure 1A). We only analyzed routes where participants successfully reached the goal (M_Goal-directed_ = 3.37, M_GPS_ = 3.16 routes). Comparing these conditions enabled us to contrast navigational signals associated with goal-directed navigation with matched motor control and optic flow, but no goal.

### fMRI acquisition and preprocessing

Participants were scanned with a 3T Siemens MRI scanner at the Rotman Research Institute at Baycrest. A high-resolution 3D MPRAGE T1-weighted pulse sequence image (160 axial slices, 1 mm thick, FOV = 256 mm) was first obtained to register functional maps against brain anatomy. Functional T2*-weighted images were acquired using echo-planar imaging (30 axial slices, 5 mm thick, TR = 2000 ms, TE = 30 ms, flip angle = 70 degrees, FOV = 200 mm). The native EPI resolution was 64 × 64 with a voxel size of 3.5mm × 3.5mm × 5.0mm. Images were first corrected for physiological motion using the Analysis of Functional NeuroImages (Cox, 1996).

All subsequent preprocessing steps were conducted using the statistical parametric mapping software SPM12 (Penny et al., 2011). Preprocessing involved slice timing correction, spatial realignment and co-registration with a resampled voxel size of 3 mm isotropic. No spatial smoothing was applied. The mean time-courses from participant-specific white matter and cerebrospinal fluid masks were regressed out of the functional images, alongside estimates of the 6 rigid body motion parameters from each EPI run. To further correct for the effects of motion which may persist despite standard processing (Power et al., 2012), an additional motion scrubbing procedure was added to the end of our preprocessing pipeline (Campbell et al., 2013). Using a conservative multivariate technique, time points that were outliers in both the six rigid-body motion parameter estimates and BOLD signal were removed, and outlying BOLD signal was replaced by interpolating across neighbouring data points. Motion scrubbing further minimizes any effects of motion-induced spikes on the BOLD signal, over and beyond standard motion regression, without leaving sharp discontinuities due to the removal of outlier volumes.

### Analysis

We used two main representational similarity analyses (Figure 1D). To maximally benefit from the temporal resolution afforded by fMRI, paths were discretized into steps: each step corresponded to a TR, or repetition time, during which an entire brain volume was measured. In the first analysis, we computed the correlation between every given step (TR) and the average of all future steps (TRs) within a particular horizon (e.g., mean of future 10 TRs following the current TR). In the second analysis, following the equations for predictive or successor representations (Dayan, 1993; Momennejad et al., 2017), we computed the correlation between every given step and the discount-weighted sum of future steps within a horizon. The pattern across voxels at each future TR was weighed exponentially using a discount parameter (i.e., gamma value, ɣ) between 0 and 1, and the value of the discount parameter corresponded to the scale of abstraction, corresponding to different levels of a representational hierarchy (Momennejad & Howard, 2018).

#### Region of Interest analysis

We investigated the predictive similarity of each state to future representations in a set of regions of interest (ROIs). To do so, we first extracted voxelwise time courses across each navigated route and z-scored the values within each voxel. We then ran two predictive similarity analyses. First, we measured the correlation of each timepoint (TR) with the mean of successor TRs within a given horizon (e.g., correlation between TR at time t, and the mean of 10 following TRs). Second, we correlated the voxelwise pattern at each timepoint (TR) within each navigated route with a discount-weighted sum of future TRs. The patterns at future TRs were weighted by different constant values (ɣ), corresponding to different predictive spatial scales. The specified ɣ values were .1, .6, .8, and .9 (Fig. 1D). With increasing ɣ values, timepoints further in the future remain weighted above zero.

As the average distance traversed within each TR was 25 meters, a ɣ value of .1 meant that only each subsequent step (1 TR away) was weighted above zero, and steps farther in the distance contributed little-to-no weight to the sum of future representations. Note that we computed the predictive horizon using the unit of fMRI measurement, i.e., a TR of 2 seconds. Hence, depending on the speed of navigation, which was matched across conditions (Fig. 2C), each step could cover a varying range of spatial distances (in meters) within and across subjects. Here we used the average distance traversed within a given horizon. For a ɣ value of .6, approximately 7 steps in the future were weighted above zero, corresponding to roughly 175 meters (Fig. 6C). For a value of .8, approximately 15 steps or 375 m were weighted above-zero, while this was the case for approximately 32 steps or 800 m for a ɣ value of .9 (Fig. 6C).

**Figure 6.**
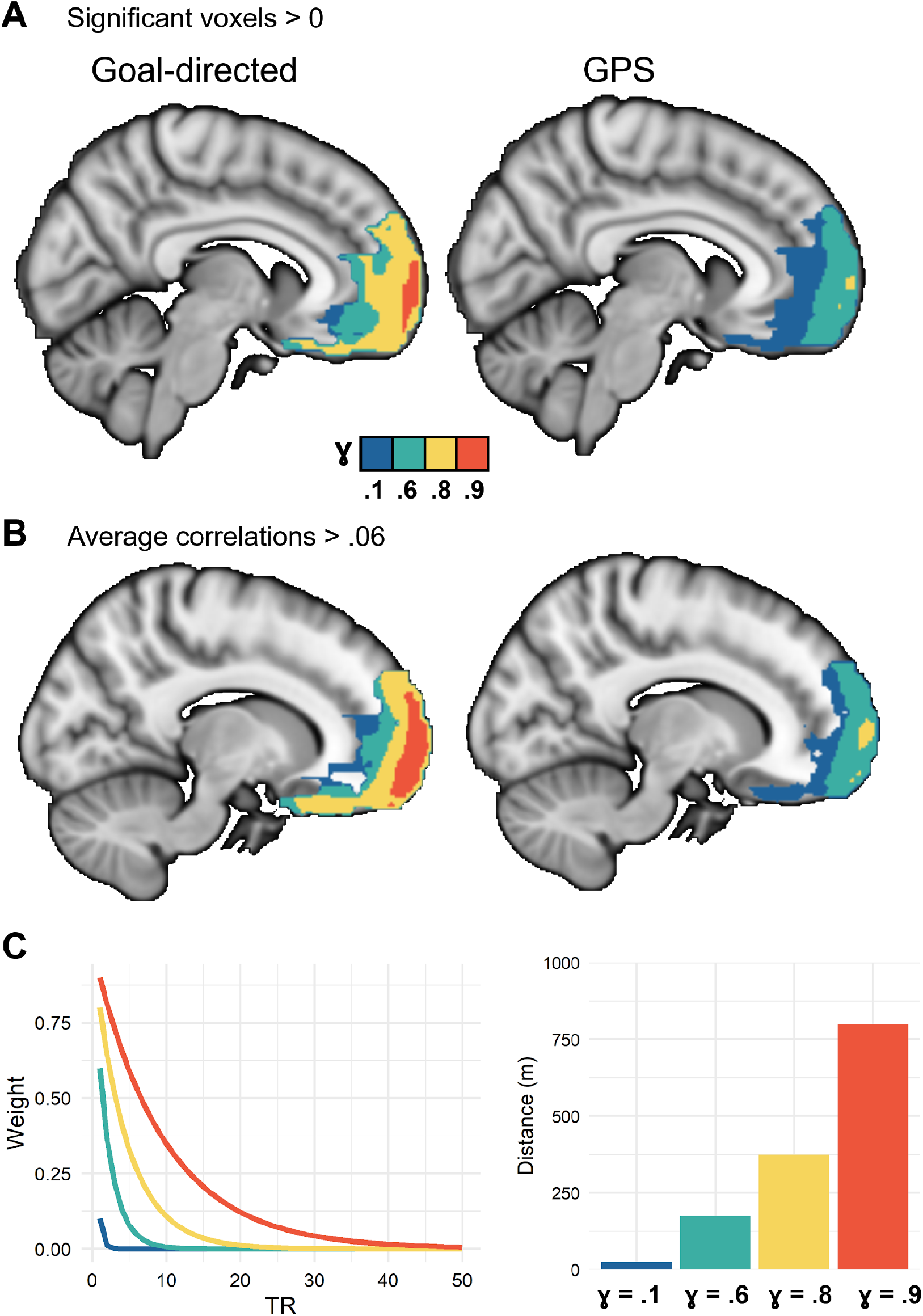
One-sample T-tests for Goal-directed and GPS condition. A) Voxels with significant representation of future states in the Goal-directed and GPS conditions using a one-sample t-test against zero. B) Voxels with representational similarity (correlation) values above .06 for each value of ɣ. *See Supplementary Figure 2 for the same results controlled for distance by adding it as a covariate within each condition*. C) Discounted weights corresponding to different gammas were applied to each successor TR. The average distance covered in each TR was approximately 25 m (24.8 m). Based on this, we computed approximate distances corresponding to predictive horizons for each discount parameter. Please note that the exact distances for each discount parameter differed across routes and participants depending on their speed. *ɣ=.1* only included 1-step (1 TR) away, *ɣ=.6* reached approximately 7 steps in the future, corresponding to roughly 175 m, *ɣ=.8*, approximately 15 steps or 375 m, *ɣ=.9* reached approximately 32 steps or 800 m ahead.

The TR-by-TR correlations within each route were averaged to derive the representation of future states on each trial. We first applied this analysis to *a priori* ROIs, including bilateral anterior and posterior hippocampi (aHPC, pHPC) and anterior and medial prefrontal cortical ROIs (antPFC, mPFC). As described in Brunec, Bellana et al. (2018), we divided the hippocampus into 6 anterior-posterior segments. We also examined the same measure in the mPFC and antPFC. The anterior PFC and medial PFC ROIs were defined as spheres surrounding peak voxels identified in preliminary findings from an fMRI adaptation of a known behavioral study of successor representations (Momennejad et al., 2017) reported in (Russek et al., 2018). The spheres were centered on an anterior prefrontal voxel (MNI coordinates × = 8, y = 68, z = 8) and a medial prefrontal voxel (MNI coordinates z = −22, y = 56, z = −10). These analyses were performed for each of the ROIs, as well as a searchlight within the prefrontal cortex.

#### Prefrontal cortex searchlight analysis

In order to identify any gradients of predictive representation in the PFC, a custom searchlight analysis was performed within a prefrontal cortex mask (created in WFU PickAtlas). The analysis was restricted to grey matter voxels, and a spherical ROI with a 6mm radius was used to iteratively correlate each TR with the discount-weighted sum of future states for voxels within each searchlight. The searchlight analysis was performed for four different values of ɣ: .1, .6., .8, and .9. The single-subject correlation maps were then compared against zero (AFNI *3dttest++*). The output z-score maps were thresholded at values corresponding to 5% false positive rates established by a cluster-size permutation simulation (AFNI *ClustSim*).

#### Model-based analysis: The discount-weighted sum of successor states

This section addresses the reasoning behind testing the successor representation hypothesis in terms of pattern similarity between a given state and the discount-weighted sum of its successor states (Figure 1). Consider an environment that consists of *n* states, some of which lead to one another. Consider *T* to be the n × n matrix of transition probabilities for one-step transitions among these n states. In a deterministic environment, when there is a transition from a given state *S_i_* to state *S_j_*, we assign 1 in the ith row and jth column of *T* (Supplementary Figure 1, left). The successor representation under a random policy can be then computed from *T* as follows (for comparison to policy-dependent SR see Momennejad, 2020):

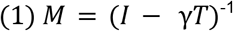

Equation (2) expands equation 1 for computing the successor representation from state *s_1_* to the goal state *s_g_* from *T,* which is one cell in the SR matrix. Recall that *T* denotes the matrix of one-step transition probabilities among adjacent states, while SR contains multi-step dependencies among non-adjacent states. Here the parameter *t* refers to the number of steps (or the distance) between states. This parameter need not denote temporal steps, and can denote any type of sequential relationship among states.

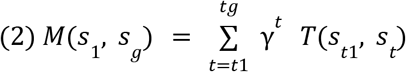

Assume the starting state is *s_1_* and the goal state is *s_5_* (as in the Markov Decision Process (MDP) in Supplementary Figure 1). Expanding equation 2, the successor representation from state 1 to 5 is *the 5th element in the 1st row of the successor representation (Equation 3), and corresponds to the expected discounted number of times we expect to visit state 5, if we start from state 1:*

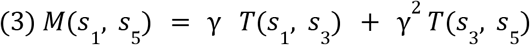

Note that equations 2 and 3 only capture 1 cell or element in the SR row associated with state *s_1_*. In the successor representation framework, the *sth* row of the SR matrix (the M matrix in Dayan 1993’s equations) is the representation we expect to observe when the agent is in state *s*. It denotes how often we expect to visit the current state’s successors on average and given a discount. A given row of the successor representation includes the present state, and the weighted representation of successor states. Thus, at the moment when an agent is in state *s,* the row activation of successor states predicts the simultaneous activation of gamma-weighted representations. We take this simultaneous row activation as the *sum of all activated weighted states in the row* (Equation 4).

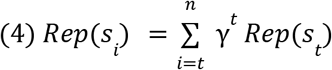

In short, the *1st row of the SR matrix* corresponds to the *representation that is simultaneously activated when the agent is in state 1, which is the sum of M*(*s_1_, s_2_*)*, M*(*s_1_, s_3_*) *, M*(*s_1_, s_4_*) *, M*(*s_1_, s_5_*). Since we only have a goal-directed trajectory, this can be the weighted sum of representations of successor states (Equation 4). Each successor state is weighted by the discount factor (gamma, *ɣ*) to the power of its distance (here in the number of states) to the starting state. A simple prediction following this weighted sum view is that being in a given state along the trajectory activates the row associated with that state and hence the weighted sum of successor states on that trajectory. This predicts neural similarity between the current state and the weighted sum of successor state representations.

Note that we did not have access to pretraining representations of the stimuli, e.g., the un-correlated representation of each location on the trajectory prior to being associated with specific paths (through lived experience in Toronto). Since we do not have these pretraining representations this method offers an approximation of the expected similarity structure. Therefore, as a general rule, we make the following prediction. In a goal-directed trajectory, and assuming the agent stays on path, we can assume that the transition probability between two adjacent states, e.g., *T*(*s_i_, s_j_*), equals 1 (i.e., we have a deterministic MDP). We predict that Equation 3 approximates the pattern similarity of the TR in the *ith* state to the weighted sum of TRs that are its successor states. Note that the predictive horizon is the successor distance within which the discount parameter γ is above zero (Figure 6). We hypothesize that different parts of the brain will show pattern similarity contingent with different values of the discount parameter 0 < γ < 1, and thus different predictive horizons.

This is a first step towards testing the multi-scale predictive representation hypothesis in a realistic navigation setting. To improve prediction accuracy, future studies are needed that incorporate diverse paths through each state, to each goal, and to different goals. These studies should include a larger graph or MDP of the environment with different starting and goal locations. In order to study map-dependent and path-dependent changes in the representation of each location, a study design is needed where the participants learn a new environment. Such studies would enable us to compare pre-training and post-training neural correlations among the states or locations in the environment.

## Results

Participants navigated a set of distances they regularly traversed in everyday life (M_Goal-directed_ = 3.5, M_GPS_ = 2.5 km). After completing each route, participants rated how familiar each route felt, and how difficult they found it to navigate on a scale from 1-9 (where 1 would correspond to least familiar and most difficult, respectively). As expected, the average reported familiarity was higher in the Goal-directed condition (M = 7.0, SD = 1.44) than in the GPS condition (M = 3.0, SD = .51; t(18) = −10.53, p < .001; Figure 2A). The subjective difficulty was similar in the Goal-directed (M = 6.98, SD = 1.43) and GPS (M = 7.2, SD = 1.08) conditions, suggesting that all navigated routes were perceived to be similarly undemanding (t(18) = .827, p = .419; Figure 2B). There was also no difference in movement speed across the Goal-directed (M = 16.21, SD = 5.04) and GPS conditions (M = 17.02, SD = 1.48; t(18) = .719, p = .481; Figure 2C). GPS routes did, however, include more turns (M = 7.08, SD = 1.39) than Goal-directed routes (M = 5.86, SD = 1.78; t(18) = 3.04, p = .007; Figure 2D). This was the case despite the GPS routes being shorter than Goal-directed routes, on average (t(18) = −4.31, p < .001; Figure 2E).

### Hippocampal and prefrontal gradients of near-future predictive representations

To investigate predictive representations along hippocampal and prefrontal hierarchies, we conducted a progression of analyses. First, we investigated representational similarity between each timepoint (TR) and the average of future *n* TRs, where *n* determined different future horizons, i.e., unweighted average of 1, 2, 3, 4, 5, or 10 future TRs (Figure 3). We conducted the analyses separately on 6 a priori regions of interest (ROIs) of anterior-posterior hippocampus, and a priori selected mPFC and antPFC ROIs (see Methods for more details). Second, we conducted the same analyses with discount-weighted sums of future TRs at different horizons (Figures 4 and 5), focusing on two posterior and anterior hippocampal ROIs and mPFC and antMPFC. In follow up analyses, we included the *path distance* on each route as a factor in the model. Third, we then conducted the discount-weighted sum RSA in a PFC-masked searchlight analysis to detect scales of representation in the PFC (Figures 6 and 7).

**Figure 7.**
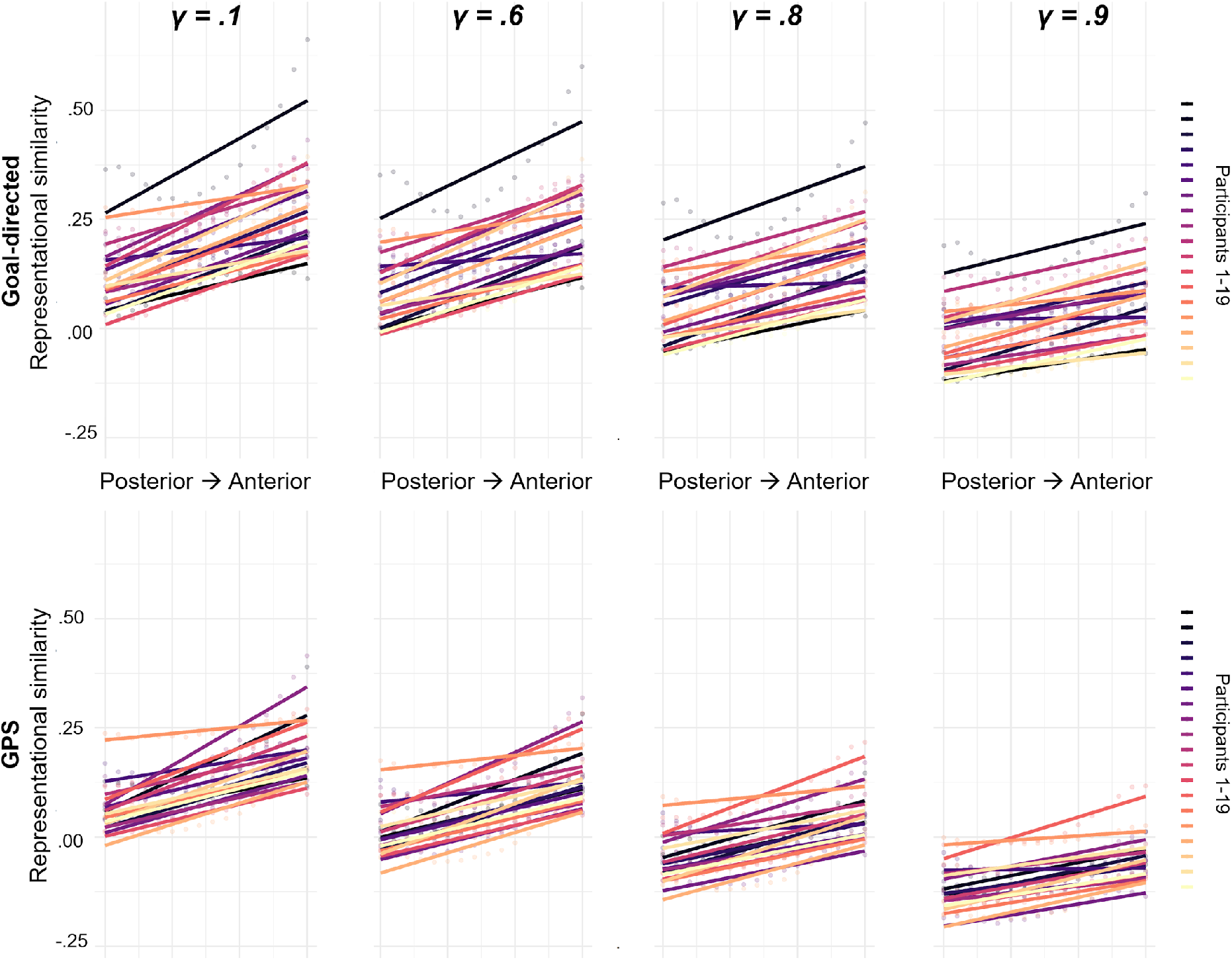
Increasing predictive similarity along posterior-to-anterior PFC. In order to indicate which PFC regions displayed higher predictive similarity we computed the slope of correlations for posterior-to-anterior PFC slices for Goal-directed and GPS conditions. We computed these slopes for four values of ɣ, corresponding to gradients of low to high scales. Each line corresponds to predictive similarity results from one of 19 participants.

### Representational similarity to mean of future TRs across horizons

We conducted linear mixed effects models on these similarity measures in bilateral hippocampi for each of the routes travelled within each condition. We included average Fisher’s z-transformed similarity on each route as the dependent variable, and axial segment (1-6), number of TRs (1-5), and hemisphere (L, R) as fixed effects. Similar analyses were performed for PFC regions of interest. Participants were included as a random effect. The random intercept mixed effects models were implemented in R (R Core Team) using the packages lme4 (Bates, Maechler, Bolker, & Walker, 2015) and lmerTest (Kuznetsova et al., 2017) to assess significance. This produced a Type III ANOVA table with Satterthwaite’s method of approximating degrees of freedom. Where these included decimal numbers, they were rounded to the nearest integer. The similarity values for 10 TRs ahead were not entered in the present model due to the non-linear shift from 5 to 10 TR, but they are plotted in Figure 3.

#### Hippocampal results

We found a significant effect of axial segment (F(5, 6796) = 45.38, p < .001), driven by greater future representations in the anterior segments compared to posterior ones. There was also a main effect of condition (F(1, 6796) = 1182.35, p < .001), reflecting generally greater values in the Goal-directed (Figure 3A), compared to the GPS condition (Figure 3B), and a significant effect of the future horizon (F(1, 6796) = 633.44, p < .001), reflecting higher similarity values for states closer to the present. There was a main effect of hemisphere, reflecting higher values in the right compared to the left hemisphere (F(1, 6796) = 6.97, p = .008). There were significant interactions between axial segment and condition (F(5, 6796) = 6.97, p < .001), axial segment and future horizon (F(20, 6796) = 2.13, p = .002), and condition and future horizon (F(4, 6796) = 13.16, p < .001). The latter interaction is of particular interest as it suggests that the decline across different predictive horizons was greater in the GPS compared to the Goal-directed condition. There was no significant three-way interaction (F < 1).

#### PFC results

We conducted the same models separately for the a priori selected regions of interest in antPFC and mPFC. In the antPFC, there was a significant main effect of condition (F(1, 1222) = 363.76, p < .001), as well as a main effect of future horizon (F(4, 1222) = 48.36, p < .001), but no condition by future horizon interaction (F(4, 1222) = 1.18, p = .319). In the mPFC, there was a significant effect of condition (F(1, 1222) = 218.77, p < .001) and a significant effect of future horizon (F(4, 1222) = 114.82, p < .001), but again no condition by future horizon interaction (F(4, 1222) = 1.82, p = .122).

Comparing the representational similarity in the Goal-directed and GPS conditions against zero, we found that the anterior PFC displayed above-zero similarity for every predictive horizon including 10 steps ahead, in the Goal-directed condition (all p-values < .001), but only up to 5 steps in the GPS condition (all p-values for 1-5 steps < .001). In contrast, the medial PFC only displayed above-zero similarity up to 5 steps in the future on Goal-directed routes (p-values < .001) and three steps on GPS routes (p-values ≤ .002). The anterior-most hippocampal segment displayed above-zero similarity for up to 4 steps in the future (p-values ≤ .006) on Goal-directed routes and only one step on GPS routes (p < .001), while the posterior-most hippocampal segment displayed above-zero similarity for one step on Goal-directed routes (p < .001), and two steps on GPS routes (p-values ≤ .006).

We next conducted similar analyses with the weighted sum of future TRs of different horizons.

### Model-based representational similarity to future TRs in ROIs

When an RL agent is in a given state during navigation, the discount-weighted sum of the successor states is the predictive representation for that state (see Methods). Therefore, we investigated the similarity between each time point and ɣ-weighted sum of the representations of future states. We ran a series of linear mixed effects models following the logic described above, including each route within each of the conditions. The models included Fisher’s z-transformed representational similarity values as the dependent variable, with ɣ and condition as fixed effects and participant as a random effect. Ɣ was modelled as an ordinal variable. For the hippocampus, the reported statistics and plotted values apply to the right hippocampus, as there was no significant difference between left and right hippocampi (all ps > .34).

#### Mixed effects analysis

The first mixed effects model included all ROIs, in order to compare average representational similarity differences across regions with different hypothesized scales. There was a significant main effect of ɣ (F(2, 1448) = 322.14, p < .001), suggesting (not surprisingly) more representational similarity within horizons that are closer to the present state. We also observed a significant main effect of condition (F(1, 1452) = 309.46, p < .001), suggesting representational similarity at higher predictive horizons in the Goal-directed compared to the GPS condition (Figure 4A–B). There was a main effect of ROI (F(3, 1448) = 547.38, p < .001), confirming the hypothesis of longer predictive horizons in the antPFC, followed by mPFC, aHPC, and pHPC. There was also a significant interaction between ɣ and condition (F(2, 1448) = 7.49, p < .001), and a significant interaction between condition and ROI (F(3, 1448) = 10.13, p < .001).

#### WIthin-ROI analyses

Follow-up mixed effects models were conducted for predictive similarity values *within* each ROI. Significance was established against a Bonferroni-adjusted value of ɑ = .0125 (for 4 ROIs). In the antPFC, there was a significant main effect of ɣ (F(2, 347) = 53.29, p < .001). There was also a significant effect of condition, with significantly higher correlations in the Goal-directed than the GPS condition (F(1, 349) = 103.42, p < .001). There was no significant ɣ × condition interaction (F < 1). In mPFC, there was again a significant main effect of ɣ (F(2, 350) = 106.39), as well as a main effect of condition (F(1, 352) = 83.19, p < .001) in the same direction as the antPFC. There was no significant *ɣ × condition* interaction (F(2, 350) = 3.44, p = .033).

In the aHPC, there was a significant main effect of ɣ (F(2, 348) = 151.90, p < .001), a main effect of condition (F(1, 350) = 128.05, p < .001), as well as a *ɣ × condition* interaction (F(2, 348) = 4.89, p = .008). As in the mPFC, this interaction reflected a steeper slope across ɣ values in the GPS condition (−.16) than in the Goal-directed condition (−.12). In the pHPC, there was a significant main effect of ɣ (F(2, 349) = 218.38, p < .001), a main effect of condition (F(1, 351) = 87.99, p < .001), and a significant ɣ × condition interaction (F(2, 349) = 3.81, p = .023), again reflecting a steeper slope in the GPS condition (−.17), compared to the Goal-directed condition (−.13).

To test for evidence of predictive representations, we conducted one-sample t-tests to test these values against zero, with an adjusted value of ɑ = .002 (24 comparisons in total). At ɣ = .1, the similarity values in all ROIs were significantly above zero in both conditions. At ɣ = .6, similarity values for all ROIs but the pHPC were significantly above zero in the Goal-directed condition. In the GPS condition, however, similarity values in neither the aHPC nor the pHPC were significantly above zero. At ɣ = .8, values in both antPFC and mPFC remained significantly above zero in the Goal-directed condition, but only antPFC remained above zero in the GPS condition. For this value of ɣ, the values in aHPC and pHPC were not significantly above zero in either condition, and were actually significantly below zero in the pHPC. This significant negative correlation could reflect the differentiation of neural patterns across time, potentially as a manner of separating experience into fine-grained units.

### Representational similarity during goal-directed navigation is related to travelled path distance

If the hippocampus and PFC represent planning processes associated with the currently navigated route, these representations should be modulated by the route path distance. To test this, we included the *path distance* on each route as a factor in the mixed effects model. Path distance was calculated as the summed change in longitude and latitude coordinates between each adjacent pair of TRs. To account for the contribution of time, we also regressed out the number of TRs on each route. The reported model-fits thus account for the variability in the amount of time spent navigating different routes. Prior to running these models, we mean-centered distances within each participant to account for different ranges travelled. We excluded 9 Goal-directed routes from a total of 8 participants, due to improbably long paths that diverged more than 1.5 km from the main path. Including these paths, however, did not change the significance of any of the results.

#### Path distance results

In the Goal-directed condition, there was a significant effect of ɣ (F(2, 630) = 186.83, p < .001), ROI (F(3, 630) = 369.49, p < .001). There was no significant main effect of path distance (F(1, 645) = 1.06, p = .304) but there were significant interactions between ROI and path distance (F(3, 630) = 38.13, p < .001) and ɣ and path distance (F(2, 630) = 6.47, p = .002; Figure 5). The plotted values in Figure 5 were estimated using the *effects* package in R (Fox, 2003; Fox & Weisberg, 2011). There was no significant interaction between Ɣ and ROI, nor a three-way interaction (both ps > .40). As predicted, we observed no main effect of path distance in the GPS condition (F < 1), nor any interactions with ROI (F < 1) or ɣ (F < 1). The main effects of ɣ (F(2, 671) = 159.40, p < .001) and ROI (F(3, 671) = 186.99, p < .001) remained significant, however.

#### ROI and path distance interactions

We conducted a linear mixed effects model for each of the ROIs, predicting representational similarity from path distance and ɣ. In the antPFC, there were significant effects of ɣ (F(2, 170) = 34.14, p < .001) and path distance (F(1, 174) = 112.14, p < .001), but no interaction between the two (F < 1). This suggests that the effect of path distance was stable across different predictive horizons in antPFC. In mPFC, the effects of ɣ (F(2, 170) = 63.77, p < .001) and path distance (F(1, 173) = 87.27, p < .001) were again significant, as was the interaction between them (F(2, 170) = 2.21, p = .113). In the aHPC, there was a significant effect of ɣ (F(2, 170) = 84.58, p < .001), a significant effect of path distance (F(1, 177) = 60.51, p < .001), and a weaker interaction between ɣ and path distance (F(2, 170) = 3.55, p = .031). Finally, in the pHPC, there were significant effects of ɣ (F(2, 169) = 156.96, p < .001), path distance (F(1, 173) = 100.15, p < .001), and a weaker interaction between the two (F(2, 169) = 3.29, p = .040).

#### Comparison to Euclidean distance

To establish how specific these results were to the traversed paths, we re-ran the models but this time included the Euclidean distance from start to goal as a predictor instead. In the Goal-directed condition, the effects of ɣ and ROI remained significant (both ps < .001), but there was no main effect of Euclidean distance (F < 1), and no significant interaction between ɣ and Euclidean distance (F(2, 631) = 1.93, p = .145). There was an interaction between ROI and Euclidean distance (F(3, 630) = 3.33, p = .019), but no three-way interaction (F < 1). In the GPS condition, the effects of ɣ and ROI were again significant (ps < .001), and there was a weaker main effect of Euclidean distance (F(1, 49) = 4.39, p = .041), but no interactions between Euclidean distance and any other factor (all ps > .50).

### Model-based representational similarity in prefrontal searchlights

Prefrontal cortex has a much larger volume than the hippocampus. In order to identify hierarchies of predictive representations comparable to hippocampal ROIs, we ran a searchlight analysis and computed similarity for voxels within every spherical searchlight (of 6mm radius). The searchlight analysis was performed for four values of ɣ (.1, .6, .8., .9) within each of the conditions. The thresholded z-score maps for different values of ɣ are displayed as overlays in Figure 6A, along with the average thresholded similarity maps within each condition (thresholded at .06; Figure 6B).

#### Prefrontal hierarchy

To capture the gradient of values from the anterior-most to the posterior-most segments of the PFC, we calculated the average value of representational similarity across voxels within each anterior-posterior slice (i.e., the y-direction). The slopes are plotted in Figure 7. These plots reveal a gradation of predictive representations extending from posterior-most to anterior-most slices of the PFC. This trend was reliable in both the Goal-directed and GPS conditions, but the representational similarity values were consistently greater in the Goal-directed condition.

To account for the proportion of different histologically-defined brain regions covered by each significant cluster, we calculated the percentage of overlap between each prefrontal Brodmann Area (BA) region and the significant voxels for each value of ɣ in each of the conditions (Table 1 and Figure 8). These percentages represent the proportion of each BA region covered by the significant thresholded clusters. We found the largest overlap between voxels in the anterior PFC (BA 10) and significant voxels in the searchlight analysis with various ɣ values. Following anterior and polar PFC was BA 11, corresponding to the orbitofrontal cortex, and then BA 25 and 32, corresponding to subgenual area or cingulate cortex and anterior cingulate cortex respectively. These regions were followed by smaller overlap in area 47, corresponding to the orbital part of the inferior frontal gyrus, areas 46 and 9 corresponding to the dorsolateral PFC, and no overlap in area 45 corresponding to the inferior frontal gyrus.

**Table 1.** Proportion of each prefrontal Brodmann Area accounted for by the significant prefrontal voxels. Results were driven from the one-sample T-test results displayed in Figure 5A. (not matched for distance). Proportions are displayed for each value of ɣ within each condition.

|  | Goal-directed |  |  |  | GPS |  |  |  |
| --- | --- | --- | --- | --- | --- | --- | --- | --- |
| | $\gamma = .1$ | $\gamma = .6$ | $\gamma = .8$ | $\gamma = .9$ | $\gamma = .1$ | $\gamma = .6$ | $\gamma = .8$ | $\gamma = .9$ |
| **BA9** | 5.8% | 5.8% | 5.6% | 0% | 5.8% | 4.3% | 0% | 0% |
| **BA10** | 59.4% | 59.3% | 54.9% | 13.9% | 59.1% | 48.0% | 4.9% | 0% |
| **BA11** | 46.5% | 44.4% | 32.6% | 6.0% | 42.6% | 21.8% | 0% | 0% |
| **BA25** | 41.1% | 37.4% | 15.9% | 0% | 21.5% | 0% | 0% | 0% |
| **BA32** | 23.9% | 21.6% | 11.2% | 0% | 21.1% | 2.8% | 0% | 0% |
| **BA47** | 16.6% | 15.6% | 5.8% | 0% | 12.7% | 1.1% | 0% | 0% |
| **BA46** | 6.8% | 6.8% | 6.8% | 0% | 6.8% | 4.2% | 0% | 0% |
| **BA45** | 0% | 0% | 0% | 0% | 0% | 0% | 0% | 0% |

**Figure 8.**
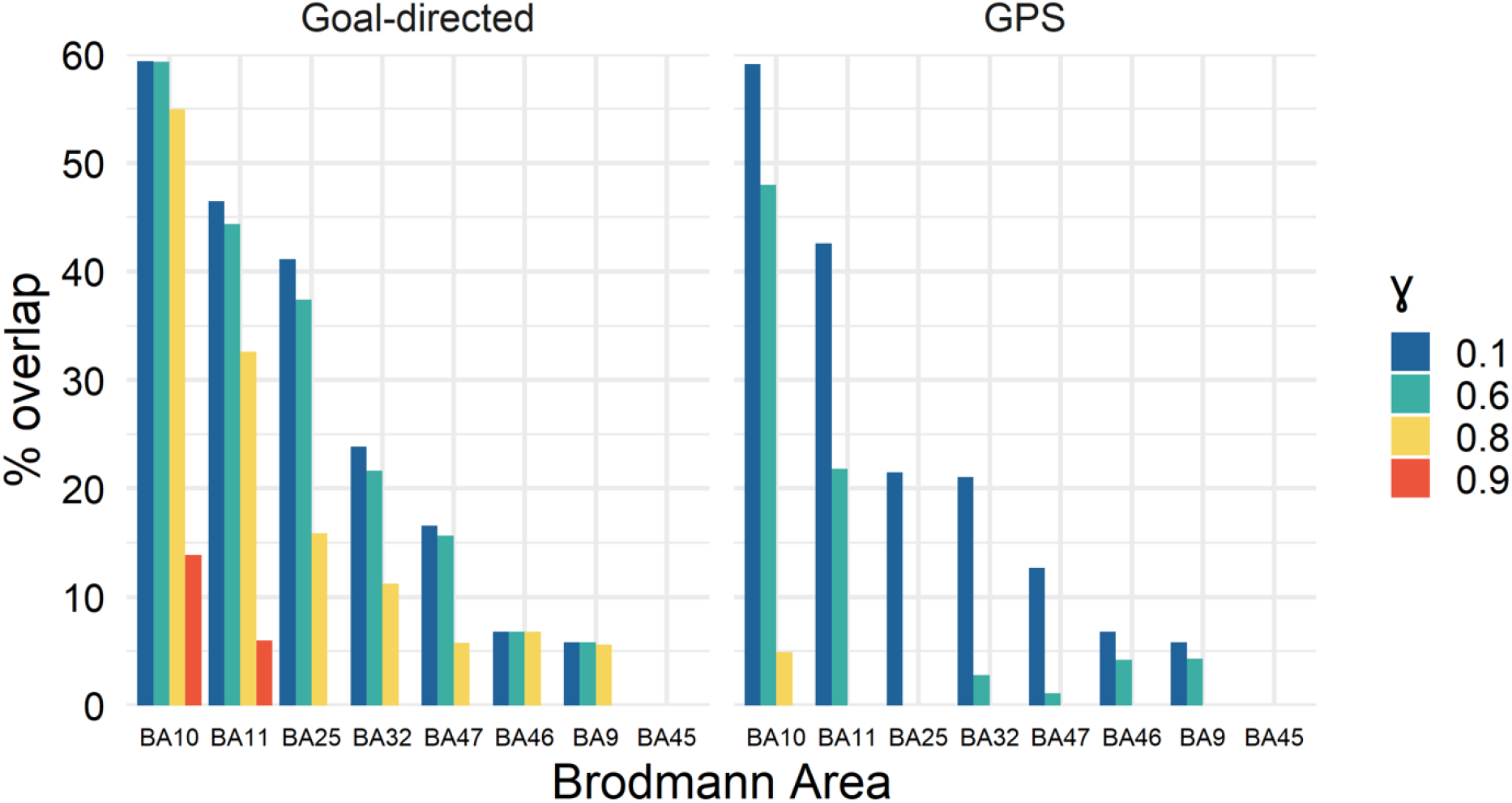
Prefrontal cortex hierarchy in the Goal-directed and GPS conditions. Proportion of prefrontal Brodmann Areas accounted for by the significant PFC voxels in searchlight analysis are shown. Results were driven from the one-sample T-test results displayed in Figure 5A (not matched for distance). Colorbars reflect different discount values (ɣ) corresponding to different predictive horizons within each condition. (Blue: ɣ=.1, Green: ɣ=.6, Yellow: ɣ=.8, Red: ɣ=.9).

### Representational similarity slope along PFC hierarchy

#### Controlling for Distance: Matched Distance Analysis

As discussed in earlier sections, the distances were not matched between the two conditions (Figure 2E). To account for this difference, we conducted a matched analysis in which we manually selected pairs of routes with the minimum difference in distance for each participant, up to a kilometer (Figure 9A). We were unable to include 3 of the participants in this analysis as the distances in their Goal-directed and GPS routes were too different (with a difference in distance > 1.5 km). For the remaining 16 participants, there was no significant difference between the selected GPS and Goal-directed routes (p = .215). We ran a paired samples t-test comparing participants’ prefrontal RSA maps for the two selected routes. We also included the difference in distance for the two selected routes as a covariate for each participant. The brain maps of the average correlation values thresholded at .04 are presented in Figure 9B and the results of the 5% FPR corrected t-test in Figure 9C.

**Figure 9.**
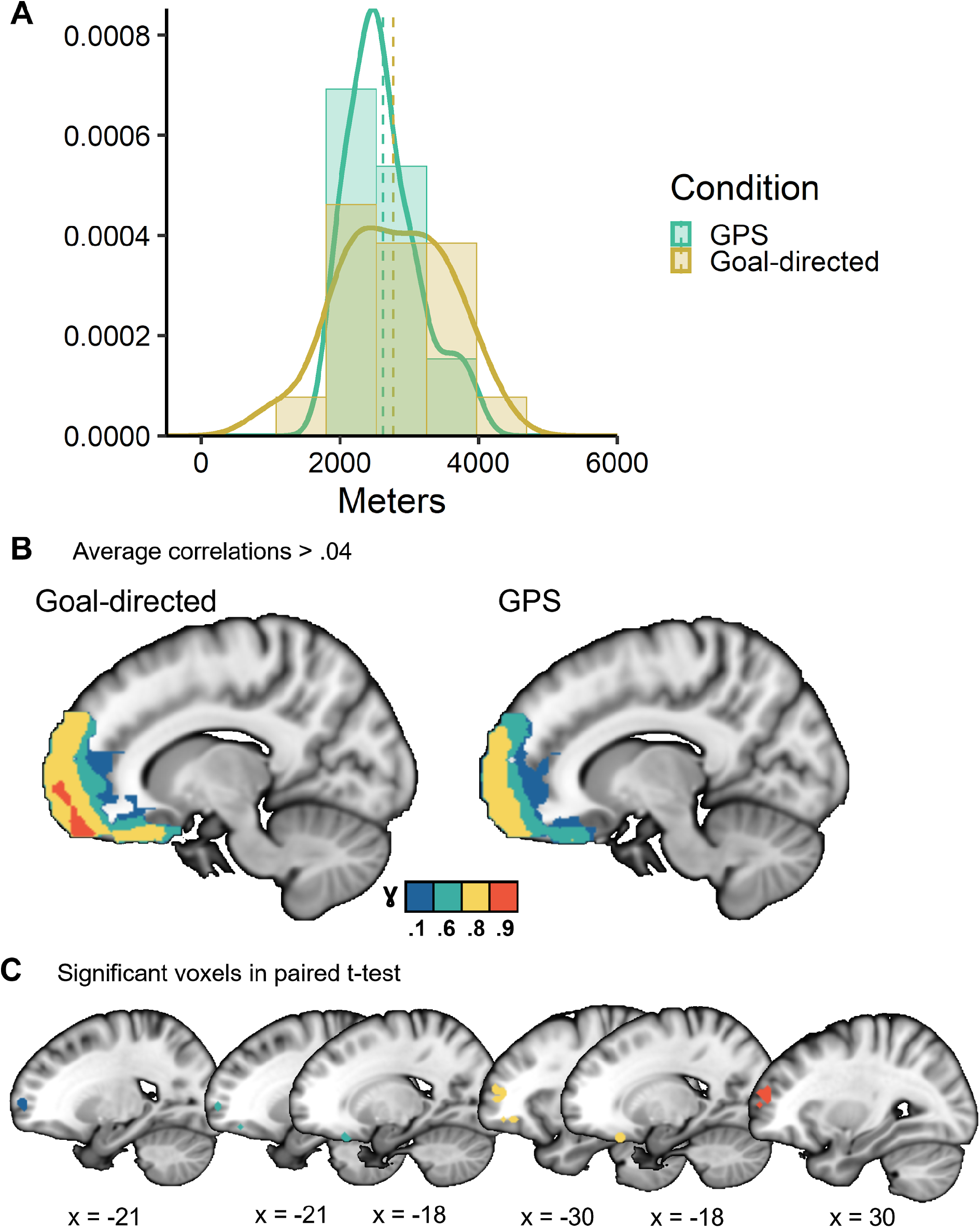
Predictive representations for Goal-directed and GPS routes with matched distances. A) Distribution of distance-matched routes included in this analysis. B) Voxels with average correlation values of > .04. C) Significant voxels in Goal-directed > GPS paired t-test, thresholded at t-value corresponding to 5% FPR. Colors reflect predictive horizons corresponding to different discount parameters (Blue: ɣ=.1, Green: ɣ=.6, Yellow: ɣ=.8, Red: ɣ=.9).

We compared matched-distance searchlight results in the Goal-directed and GPS conditions. In this comparison, relatively few clusters significantly differed between the Goal-directed and GPS conditions. However, the comparison at each level of ɣ suggests that there is a set of clusters along the rostrocaudal extent of the PFC which differentiates between goal-directed and GPS-guided navigation (Table 1). Notably, while only orbitofrontal clusters were significantly different for smaller horizons, more dorsal and rostral/polar PFC clusters emerged in the comparison of larger horizons or scales–between the Goal-directed and GPS conditions. It is worth noting, however, to ensure matched distances between the Goal-directed and GPS condition we excluded individuals with a large difference between the distances in the two conditions. As a result, this analysis only included individual paths from 16 participants, which likely results in increased noise and lower statistical power, which is common when using more naturalistic data.

## Discussion

We investigated the hypothesis that relational knowledge – about navigational paths – is organized as multiscale predictive representations in hippocampal and prefrontal hierarchies. We found evidence for such multiscale representations in a task where participants navigated the city of Toronto virtually in both Goal-directed and GPS-guided conditions, with realistically long distances (average 3 km). Our fMRI representational similarity analysis between each state (TR) and a discounted sum of its prospective states (at multiple scales, 25-875 meters) confirmed this hypothesis. These results support the hypothesis that prefrontal-hippocampal representations organize relational knowledge–in this case for spatial navigation–at different scales of generalization and abstraction (Behrens et al., 2018; Momennejad & Howard, 2018).

We have reported four main findings. First, as predicted, fMRI similarity reflected longer predictive horizons for paths in the Goal-directed, compared to the GPS condition. Second, similarity values in the anterior hippocampus and anterior PFC were significantly higher in the Goal-directed condition and for longer horizons. Third, predictive representations were organized along a posterior-anterior hippocampal gradient of predictive horizons (25-175m) with larger scales in gradually more anterior hippocampal regions. Fourth, representational similarity to future horizons was organized along a rostro-caudal gradient in the PFC with larger-scale horizons (25-875m) in gradually more anterior regions (Figure 6).

### Representational hierarchy

In spatial navigation, this hierarchical representation could enable hierarchical planning and subgoal computation (Ribas-Fernandes et al., 2018). Our proposal is that planning at larger scales may be enabled by larger and more abstract scales of predictive representations in anterior PFC (Figure 1, large scale graph). This higher level plan may be translated into more precise policies using representations in pre-polar PFC, orbitofrontal cortex, and anterior hippocampal regions (Figure 1, mid-scale graph), and finer scale trajectories are translated by hippocampal gradients down to the smallest predictive horizons of place fields (Figure 1, small scale graph).

### Hippocampal hierarchy

These multiple scales of representation can be simultaneously activated, while the spatial precision of information differs across the scales. It is possible that the global structure of each route is thus represented in the PFC, while more fine-grained representation of individual locations is supported by the hippocampus. This is consistent with recent work on cognitive maps in rodents, monkeys, and humans indicating PFC’s involvement in *active* navigation and planning (Epstein et al., 2017), as well as previous work suggesting that the place fields are gradually larger along the dorsal-ventral axis of the hippocampus (Strange et al., 2014), and that long axis of the hippocampus supports gradually larger spatiotemporal scales (Brunec et al., 2018; Nielson et al., 2015; Poppenk et al., 2013; Strange et al., 2014; Peer et al., 2019) and inference on mnemonic relations (Collin et al., 2015; Schlichting & Preston, 2015). Recent computational perspectives suggest that the hippocampus and PFC form and update a predictive map of the state space at multiple scales (Momennejad & Howard, 2018), organizing relational knowledge of spatial and non-spatial states (Garvert et al., 2017; McKenzie et al., 2014; Schuck et al., 2016; Bellmund et al., 2018).

### Nonspatial relevance

We have previously proposed a role for hierarchies of predictive representations along prefrontal and hippocampal gradients (Momennejad & Howard, 2018). Similar representational hierarchy may also underlie relational knowledge and category generalization (Constantinescu et al., 2016), abstraction and transfer (Cole et al., 2011), reward predictions (Takahashi et al., 2017), associative inference, and schema learning (Hebscher & Gilboa, 2016; Moscovitch & Melo, 1997; Spalding et al., 2018; van Kesteren et al., 2013; Yu, 2018; Zeithamova et al., 2012; Zeithamova & Preston, 2010). Previous work has proposed a hierarchy of time-scales in the brain (Chen et al., 2015) and indicated a role for hippocampal-prefrontal interactions in integrating episodes to build abstract schema (Schlichting & Preston, 2017).

### Prefrontal hierarchy

Comparing predictive similarity values across the PFC gradient (Figures 6-9, Table 1) revealed an overall effect of condition and prefrontal gradient. Longer predictive similarity horizons were observed in the Goal-directed vs. the GPS-guided condition, and more anterior PFC regions showed predictive similarity for longer horizons. Furthermore, we measured the percentage of overlap between significant voxels in the searchlight analysis and voxels in different histologically-defined PFC regions. We found the largest overlap in the anterior PFC (BA 10), orbitofrontal cortex (BA 11), and granular and anterior cingulate cortex (BA 25 and 32)–consistent with the direction of the slope of predictive similarity in Figure 7.

### The anterior PFC

The majority of PFC voxels that displayed larger predictive similarity were in polar or anterior PFC (BA10, Table 1, Figure 8). BA 10 is the largest cytoarchitectonic region of the human PFC, it has the largest volumetric and proportional difference between humans and other great apes, it is highly interconnected within the PFC, and its cells display longer decay times (Ramnani & Owen, 2004). Thus, the properties of BA 10 suggest a structurally well-connected region to support higher scales of abstraction. This includes supporting predictive representations with larger scales of integration, which can be thought of in terms of clustering of relational graphs with a higher radius. For temporal relations, this graph clustering or integration radius can be thought of in terms of longer decays or longer sustained memory leading to binding over longer time-scales. For spatial relations, this radius can be thought of in terms of associating locations that are farther apart. For relational structures, this radius can be thought of in terms of an increase in similarity among a cluster of associations within a given degree of separation. Such a prefrontal hierarchy of relational abstraction could therefore support task sets and schema.

### Relation to task sets and prospective memory

A crucially non-spatial body of evidence from the study of goal-directed tasks and prospective memory is relevant here. These studies indicate a functional role for anterior or rostral prefrontal cortex in the encoding and retrieval of prospective task sets and goals (Gilbert, 2011; Haynes & Rees, 2006; Momennejad & Haynes, 2012, 2013). This frontopolar evidence fits well with the proposal that the PFC is organized in a rostrocaudal hierarchy (Badre & D’Esposito, 2007; Koechlin, 2011; Koechlin et al., 2003; Koechlin & Hyafil, 2007), with more anterior or rostral regions corresponding to higher levels of integration and relational abstraction (Bunge et al., 2003; Kalina Christoff et al., 2009; Momennejad & Haynes, 2013). Lesions to the frontopolar cortex do not impair usual navigation or performance on intelligence or working memory tests, but impair the patient’s ability for multi-tasking and prospective memory (Burgess, 2000; Volle, Gonen-Yaacovi, Costello, Gilbert, & Burgess, 2011) such as completing a sequential plan for simple everyday tasks, e.g., plan a visit to multiple stores on a street to write a note and stamp and post it (Burgess, 2000).

### The orbitofrontal cortex

We also found that more anterior orbitofrontal cortex (OFC) yielded higher predictive similarity along larger predictive horizons, more similar to aPFC than the hippocampus (Figures 4, 8). In fact, OFC and anterior PFC have both been suggested to support model-based reinforcement learning (Daw et al., 2011), where an animal unfolds a learned state-action-state associative model during goal-directed planning and decision-making. This finding has been replicated across different experiments (Daw & Dayan, 2014; McDannald et al., 2012, 2014; Pauli et al., 2019). It supports the idea that the OFC maintains task-relevant state-state relational maps that enable iterative value computation in planning and decision-making (Daw et al., 2005; Keiflin et al., 2013; Simon & Daw, 2011; Wilson et al., 2014). We also found that predictive representations in the anterior hippocampus were the most similar to OFC representations. This is consistent with recent work on OFC-hippocampal interactions in model-based behavior (Miller et al., 2017; Vikbladh et al., 2019; Keiflin et al., 2013; Schuck et al., 2016; Wikenheiser & Schoenbaum, 2016; Wood & Grafman, 2003).

Note that prefrontal hierarchies of representations need not be static. These representations could be constructed from compressed representation, e.g., eigenvectors (Stachenfeld et al., 2017), inverse laplace transform (Ida Momennejad & Howard, 2018) (Momennejad, Howard, 2018), or generative models (Whittington et al., 2019). Future studies are required to shed light on prefrontal and medial temporal contributions to the process involved in integration, eigen-decomposition, generative models, and abstraction.

### Caveats and future direction

The fMRI dataset used here (Brunec, Bellana et al., 2018) was acquired as participants moved through long distances of 1-5 kilometers in a virtual rendering of a city they lived in (Toronto). However, the data set has some caveats, some of which we addressed in our controlled analyses, and some of which remain to be more directly addressed by future studies.

The first caveat was that the navigated routes in the Goal-directed condition were, on average, significantly longer than those in the GPS-guided condition (Figure 2). To overcome this caveat, we first controlled for distance in one-sample t-tests to reveal regions with significant pattern similarity within a given horizon (Supplementary Figure 2). In a more conservative analysis, we reran the analyses excluding longer routes and including only Goal-directed routes that were within the range of distances in the GPS-guided condition (Figure 9). Consistent with our earlier findings, longer predictive scales engaged more dorsal PFC regions significantly more in the Goal-directed condition. These controlled analyses suggest that our main findings are reliable (c.f., Table 1 and Figures 7 and 8). However, future studies with a controlled design of traversed distances and more participants are needed to replicate these findings.

The second caveat is that the selection of routes did not include multiple past and future trajectories for each state (especially subgoals), nor multiple past routes for each goal location. Future studies with such a design would allow for testing of the graph structure of relations. Such a study would also advance previous work using routes with multiple paths (Balaguer et al., 2016; Chanales et al., 2017), and allow us to dissociate pattern similarity due to the *memory of the past* from pattern similarity due to *predictive representations* more directly.

Measuring neural representations as participants navigate a full graph would also enable investigating compressed representations. For instance, we could study whether states (e.g., subgoals, and other states with special graph properties) that appear on many paths have a pronounced predictive representation, or whether nearby locations are clustered as one state (or subgoal) by some brain regions. Such a design could be implemented in future fMRI studies, enabling a more thorough analysis of prefrontal-hippocampal interactions in abstraction and subgoal processing during planning.

Finally, future studies could test the temporal hierarchy of large-scale predictive representations in the PFC and the hippocampus for higher level plans (e.g., train from New York to Philadelphia) and smaller subgoal processing (e.g., walk to the train station first). They can also test the dynamics of goal and subgoal representation more closely, complementing existing electrophysiology and neuroimaging work showing goal representation in the hippocampus (Brown et al., 2016; Howard et al., 2014; Sarel et al., 2017; Tsitsiklis et al., 2019).

### Other interpretations

Theoretically, we have proposed that a given state has higher representational similarity to its frequently visited successors along the planned path (Ezzyat & Davachi, 2014; Garvert et al., 2017; Momennejad et al., 2017; Stachenfeld et al., 2017). Our interpretation can also be discussed in terms of increased association, integration, abstraction, and clustering (Ritvo, Turk-Browne, & Norman, 2019), or the spread of activation across memory networks (Sievers & Momennejad, 2019). One may also ask whether the similarity to successor states reported here could be due to the replay of previous trajectories or paths (Ambrose, Pfeiffer, & Foster, 2016; Momennejad et al., 2018; Wu & Foster, 2014). While there are clever analytic designs to hint one way or another, a clear-cut dissociation of these hypotheses requires higher spatio-temporal resolutions such as electrophysiology, MEG, and other methods across species.

### Summary

We present support for the hypothesis that predictive maps with different scales are structured in hippocampal-prefrontal hierarchies. We found that while posterior hippocampal regions supported smaller predictive scales–up to 100-200 meters–anterior prefrontal regions supported larger predictive horizons–which in this spatial navigation task extended to 875-900 meters. Our results support the idea that medial temporal-prefrontal representations underlie cognitive maps and hierarchical planning. The organizational principles of predictive hierarchies can be extended to non-spatial domains. Future studies can be specifically designed to further investigate planning, subgoal setting, and abstraction in spatial and non-spatial graphs.

## Acknowledgements

The authors gratefully acknowledge Morris Moscovitch and Jason Ozubko for designing the original experiment, collecting and sharing the data, and sincerely thank Ken Norman, Buddhika Bellana, and Morgan Barense for helpful discussions. This work was supported by Canadian Institute of Health Research grant (#MOP49566 and #MOP125958) to Morris Moscovitch, a doctoral award from the Alzheimer Society of Canada (I.K.B.), grants from the James S. McDonnell Foundation and Natural Sciences and Engineering Research Council granted to Morgan Barense, NIMH Grant R01-MH104606 granted to Joshua Jacobs, and the John Templeton Foundation (I.M.), NIBIB R01EB022864.

## Supplementary Material

### Supplementary Methods

**Supplementary Figure 1.**
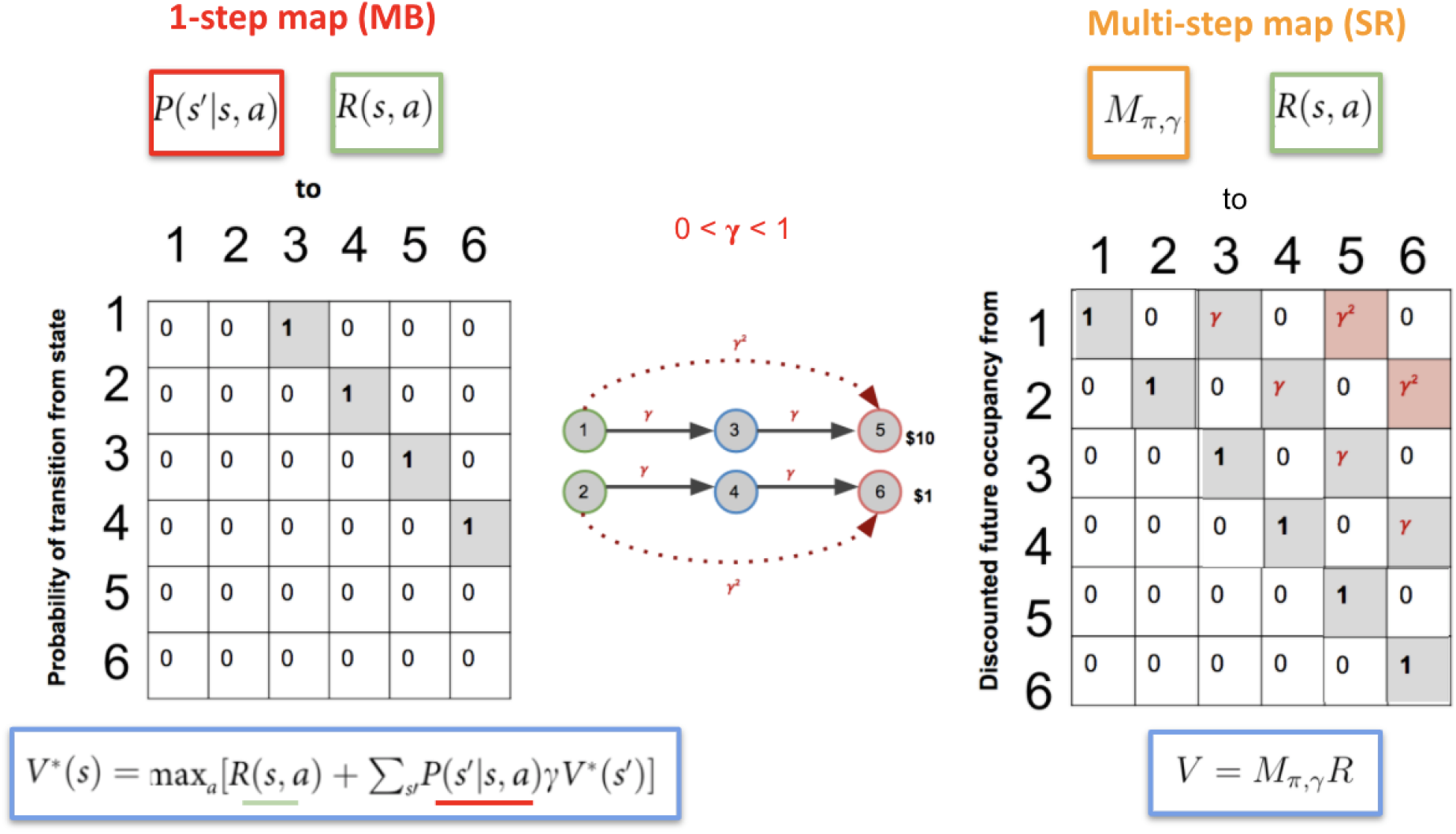
Assume an environment with the structure in the graph or Markov Decision Process (MDP) below. The starting state is *s_1_* and the goal state is *s_5_* (as in the MDP below), where we observe the highest reward. On the left we have the 6×6 transition probability matrix T, with probability 1 for every 1-step connection between any 2 adjacent states. On the right we have the successor representation matrix, which can be computed As follows:

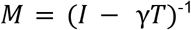

Note that the *1st row of the SR matrix* corresponds to the *representation when the agent is in state 1*. Consider an environment with 6 states and relationships depicted in the Markov Decision Process (MDP) in Supplementary Figure 1. In reinforcement learning, computing the value (*V*) of a state, under policy π, can be arrived at by multiplying the associated SR row by the vector of rewards (a 6 item vector storing the observed reward for each state).

**Supplementary Figure 2.**
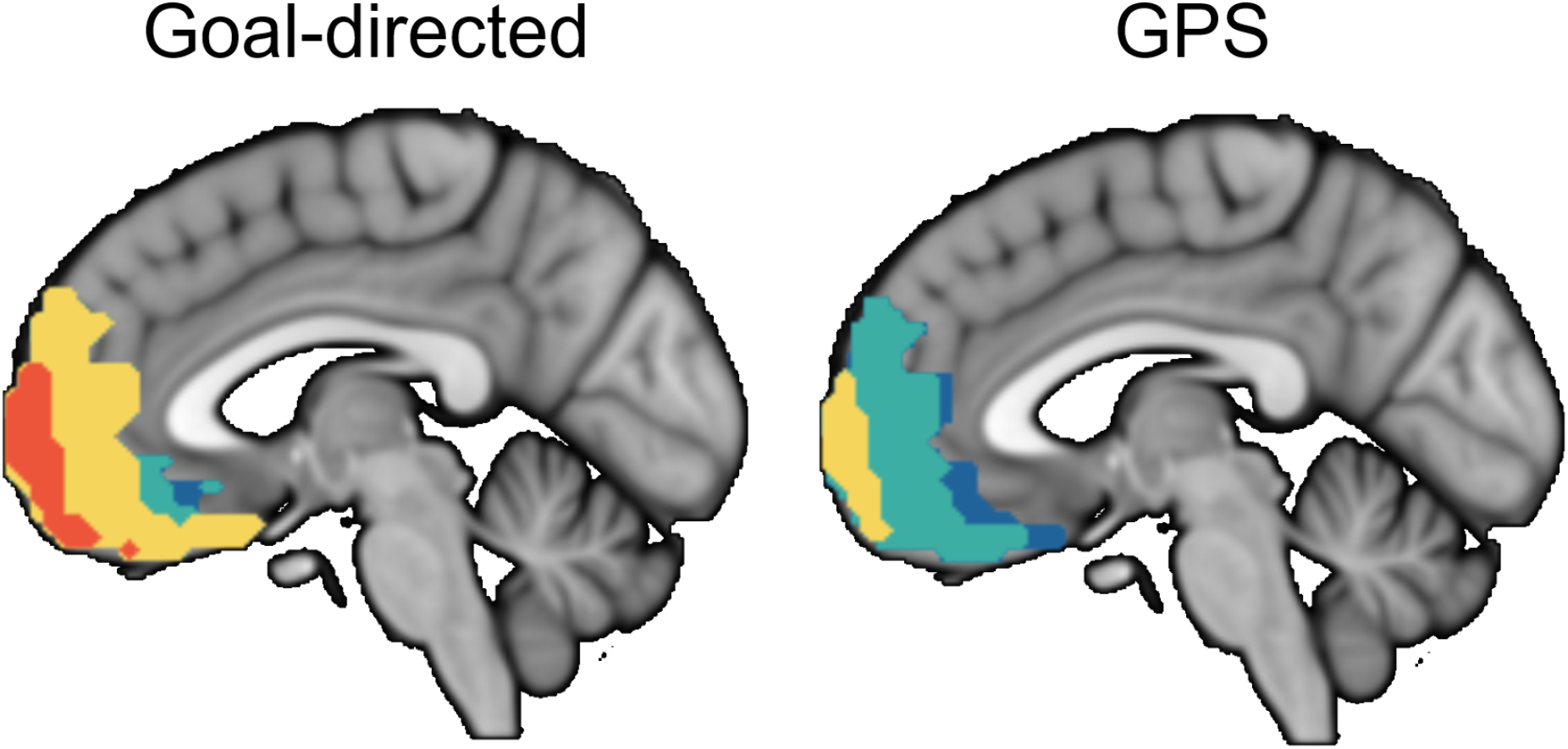
One-sample T-tests with distance as covariate. The results look very similar to running to running a T-test on Goal-directed routes vs. zero, and GPS routes vs. zero. The mean distance, per participant, per condition, was included as a covariate.

